# HiCAT: A tool for automatic annotation of centromere structure

**DOI:** 10.1101/2022.08.07.502881

**Authors:** Shenghan Gao, Xiaofei Yang, Xixi Zhao, Bo Wang, Kai Ye

## Abstract

Significant improvements in long-read sequencing technologies have unlocked complex genomic areas, such as centromeres, in the genome and introduced the centromere annotation problem. Currently, centromeres are annotated in a semi-manual way. Here, we propose HiCAT, a generalizable automatic centromere annotation tool, based on hierarchical tandem repeat mining and maximization of tandem repeat coverage to facilitate decoding of centromere architecture. We applied HiCAT to human CHM13-T2T and gapless *Arabidopsis thaliana* genomes. Our results not only were generally consistent with previous inferences but also greatly improved annotation continuity and revealed additional fine structures, demonstrating HiCAT’s performance and general applicability.

## Background

Centromeres play an essential role in the transmission of genetic information between generations. Deep analysis of centromere architecture is critical to understanding genome stability, cell division and disease development[1]. In most eukaryotes, centromeres exhibit extra-long tandem repeat (TR) sequences, but the sequence and length of repeat units, which are referred to as monomers, vary significantly among species[2]. The canonical order of monomers yields higher order repeats (HORs)[3]. For example, in the active centromere of the human X chromosome (CENX), 12 monomers (the length of one monomer is approximately 171 bp) are consecutively ordered as HOR units (the length of one HOR unit is approximately 12×171 bp) (Fig. 1a)[4]. The sequence identity between monomers within an HOR unit is only 50–90%, but the pairwise sequence identity between HOR units in a given centromere is as high as 95-100%[5]. The extra-long TRs and high homogeneity make it difficult to achieve accurate assembly of centromeres, hindering thorough investigations of their sequence architecture[5]. The rapid development of long read sequencing technologies, especially PacBio high-fidelity (HiFi) reads, has greatly improved genome assembly quality[6]. Based on this progress, the Telomere-to-Telomere (T2T) consortium presented the complete sequence of the human complete hydatidiform mole (CHM) cell line CHM13 in 2022[7]. In addition, gap-free genome assembly has been achieved in a few plant genomes, such as those of *Arabidopsis thaliana* and *Oryza sativa*[8, 9]. Significant improvements in genome quality have also contributed to the development of bioinformatic methods for the study of centromere architecture.

**Fig. 1.**
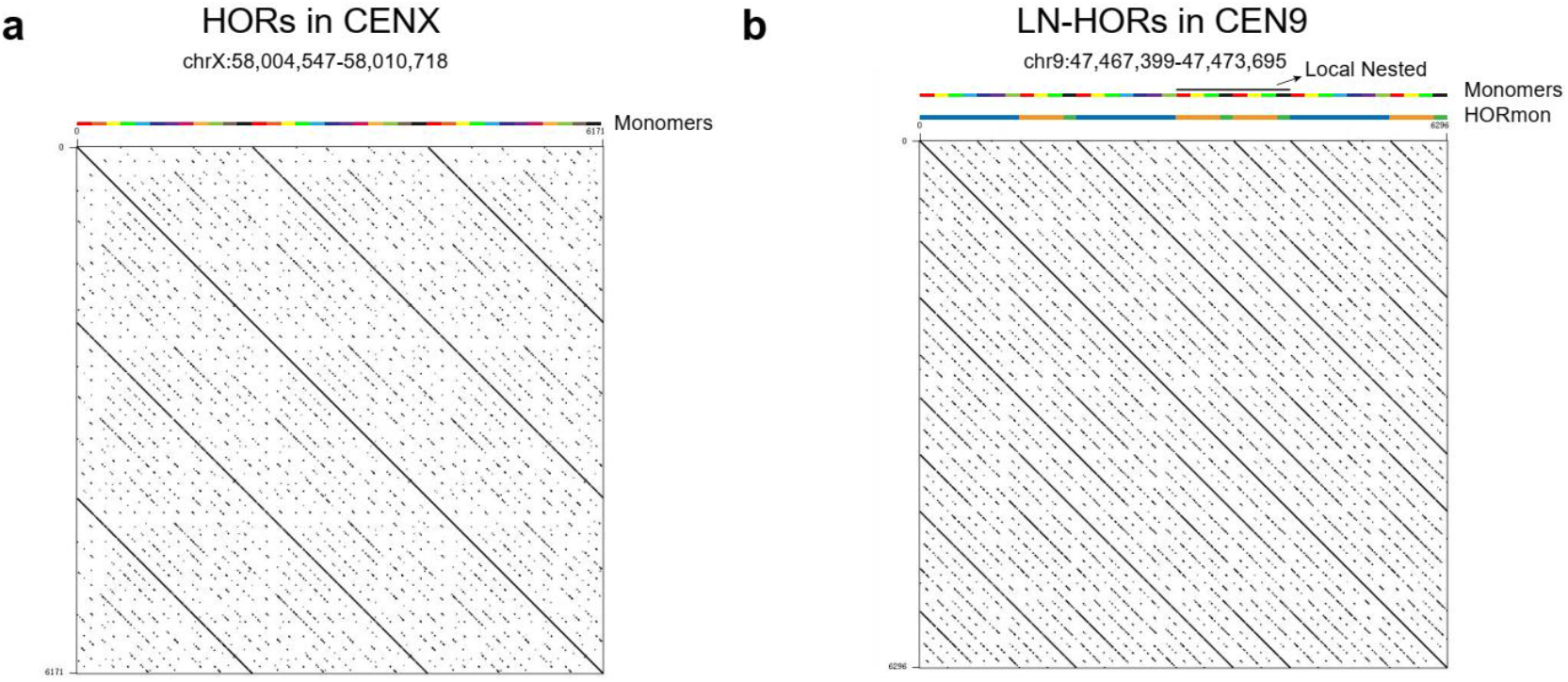
Examples of higher-order repeats (HORs). **a**. HORs in CHM13 CENX. **b**. Local nested HORs (LN-HORs) in CHM13 CEN9. In the monomer tracks, rectangles in various colours represent different monomers. In the HORmon tracks, differently coloured rectangles represent different annotations in HORmon. Blue, orange and green rectangles represent the annotated canonical HORs, partial HORs and monomers not belonging to any HORs, respectively.

Centromere annotation, including monomer inference and HOR detection, is a prerequisite for studying the structure and evolution of centromeres within and between species[10]. Previous studies annotated a substantial number of monomers and HORs in the human genome in a semi-manual manner, facilitating the understanding of centromere architecture[11-13]. However, this semi-manual method lacks a rigorous algorithm definition and is time-consuming and laborious, prohibiting its ready application to new assemblies. To address this question, Dvorkina et al. proposed the first automatic centromere annotation tool, CentromereArchitect[10], which was based on StringDecomposer (SD)[4], an algorithm for detecting sequence blocks by taking monomer templates to decompose centromere DNA sequences. In CentromereArchitect, monomer inference and HOR detection were considered two separate problems without interconnections, which often led to biologically inadequate annotation[14]. The authors next proposed HORmon[14] based on the centromere evolution postulate (CE postulate, where each monomer appears only once in the HOR unit) to address the lack of interconnection issue in CentromereArchitect. HORmon first constructs a *de* Bruijn graph based on monomers inferred from CentromereArchitect and then refines the monomers by considering positional similarity to amend the graph as a single cycle (referred to as the detected HOR) to comply with the CE postulate. Finally, HORmon classifies the detected HORs into canonical and partial HORs. However, the CE postulate has never been strictly proven and heavily depends on parameters[14], while a single occurrence of each monomer in a HOR does not always hold. For example, TR expansion does occur within HORs and forms so-called local nested HORs (LN-HORs) (Fig. 1b). Specifically, human CHM13 CEN9, 13 and 18 have various lengths of HOR units within each chromosome, and these HORs contain shared monomers (Additional file 1: Fig. S1) due to local nesting, violating the CE postulate[14]. Thus, a substantial number of partial HORs were introduced based on the CE postulate, breaking annotation continuity and hindering the characterization of fine internal architectures in these centromeres (Fig. 1b). To overcome these problems, we propose a generalizable automatic centromere annotation tool named HiCAT based on hierarchical tandem repeat mining (HTRM) using a bottom-up iterative TR compression strategy to detect and represent LN-HORs, achieving **Hi**erarchical **C**entromere structure **A**nno**T**ation. In addition, by maximizing TR coverage, HiCAT automates parameter selection and optimizes both monomer inference and HOR detection simultaneously. We applied HiCAT to newly assembled telomere to telomere (T2T) genomes of human[11] and *Arabidopsis thaliana*[8]. We compared the results from HiCAT and those from semi-manual and HORmon approaches. We found that our automated results are generally consistent with those of previous studies. In addition, HiCAT greatly improved annotation continuity and was able to detect fine structures that were missed by other methods. All the comparison results demonstrate the superior performance and generalization of HiCAT.

## Results

### Overview of HiCAT

HiCAT takes a monomer template and a centromere DNA sequence as inputs. There are two steps in HiCAT: generation of a block list and similarity matrix (Fig. 2a) and mining of HORs (Fig. 2b). In the first step, HiCAT uses StringDecomposer[4] to transform a centromere DNA sequence into a block list based on an input monomer template. Each block is a subsequence of the centromere DNA sequence and exhibits high similarity to the monomer template. Then, we defined a similarity score based on the block edit distance to obtain a block similarity matrix (Methods). To improve calculation efficiency, we pre-processed the block similarity matrix by merging identical blocks. In the second step, to optimize monomer inference and HOR detection at the same time, we applied a TR coverage maximization strategy to guide parameter selection and establish feedback between monomer inference and HOR detection. We defined a block graph whose nodes are blocks and edges are links between any two blocks if their similarity value is greater than a given similarity threshold. A series of graphs are created when the similarity threshold iteratively increases from the minimum value (by default 94%) to nearly 100% with a specific step (by default 0.5%). For each constructed block graph, we used the Louvain algorithm[15, 16] to detect block communities, i.e., so-called monomers. We assigned a unique number to each detected monomer as its ID and transformed the block list into a monomer sequence. To detect LN-HORs, we proposed the hierarchical tandem repeat mining (HTRM) method (Methods, Additional file 1: Fig. S2 and Additional file 1: Supplementary method). HTRM recursively detected and compressed local TRs in the monomer sequence until no TRs were identified. After HTRM, we merged all TRs with shifted monomer pattern units, such as 1-2-3-4, 4-1-2-3, 3-4-1-2 and 2-3-4-1, to obtain HORs. We calculated the associated HOR coverage of each similarity threshold and chose the threshold with the largest coverage to obtain HiCAT HORs. Finally, we scored HORs based on coverage and the degree of local nesting to rank all HORs (Methods). Each HOR was named “R + (rank) + L + (length of HOR unit in the monomer pattern)”. For example, the first HOR in human CENX with 12 monomers was named R1L12.

**Fig. 2.**
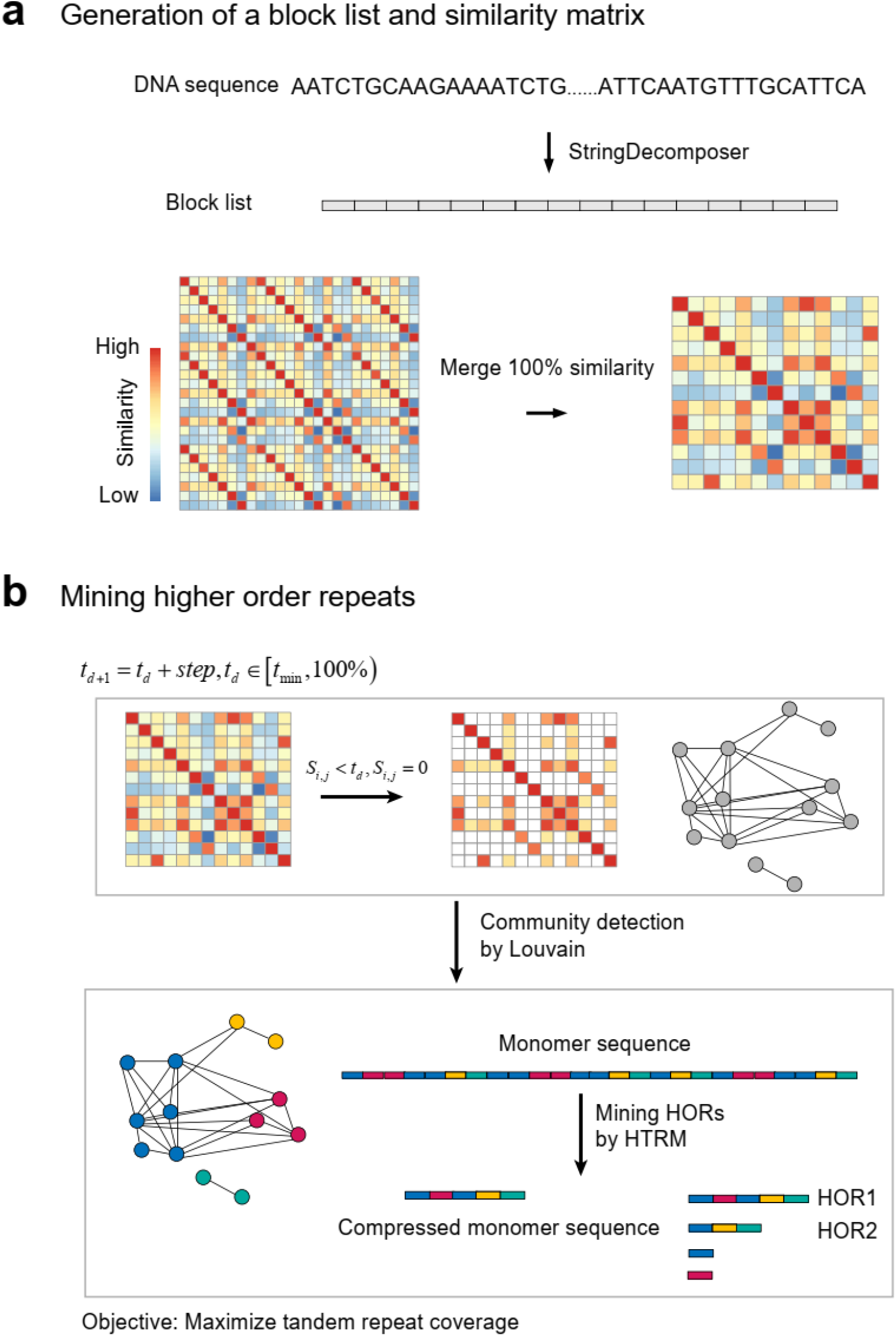
Overview of HiCAT. **a**. Generation of the block list and similarity matrix. **b**. Mining of higher order repeats (HORs). *t*_*d*_ represents the similarity threshold in the current iteration. *t*_*d* +1_ represents the similarity threshold in the next iteration. *t*_min_ is the minimum similarity threshold. *step* is the threshold increase for each iteration. *S*_*i, j*_ is the similarity between block *i* and block *j*. HTRM: hierarchical tandem repeat mining. Coloured rectangles in the monomer sequence represent monomers.

### Overall performance for human CHM13 centromeres

We first applied HiCAT in an active alpha satellite array for each centromere (Additional file 2: Table S1) of the human CHM13-T2T genome (v1.0)[11] and compared the results with published results obtained with semi-manual inference[11, 13]. We found that the HiCAT results were highly consistent with those of previous studies. The reported HORs in 21 out of 23 centromeres (CEN1, 2, 3, 4, 6, 7, 8, 9, 10, 11, 12, 13, 14, 15, 16, 18, 19, 20, 21, 22 and X) were well detected by HiCAT, while substantial differences were observed for the remaining two chromosomes, CEN5 and CEN17 (Additional file 3: Table S2). We first took CEN11 and 15 as examples to further explore the HiCAT results. There were two types of HOR units, nested (LN-HORs) and canonical. We found that HORs in CEN11 were rather homogeneous, with as few as 12 nested units in R1L5 (Fig. 3a, b). In CEN15, there were approximately four times as many nested units in R1L11 as canonical units (Fig. 3c, d). The monomer pattern of the CEN15 R1L11 unit was (6-7-14-10)×n-1-2-8-3-9-11-4. Each number represents a monomer, and “×n” represents the number of times that a defined monomer set was repeated. For example, four consecutive monomers 6-7-14-10 in the R1L11 unit experienced expansion, and most of them expanded twice, while other numbers of repeats also existed (Fig. 3e).

**Fig. 3.**
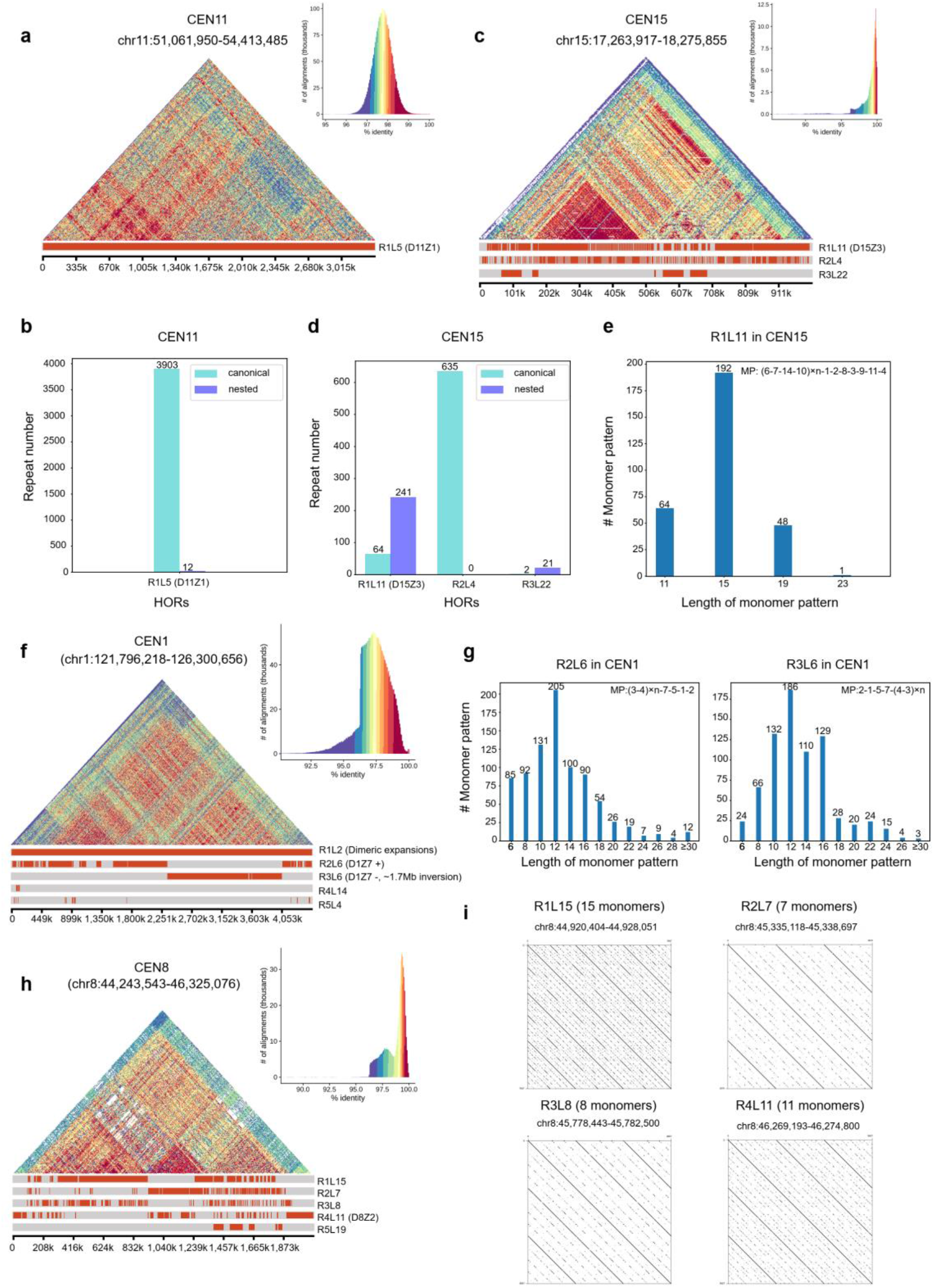
Fine structures in CHM13 CEN11, 15, 1 and 8. **a**. Structure and annotation of CEN11. **b**. The numbers of HOR repeats in CEN11. **c**. Structure and annotation of CEN15. **d**. The numbers of HOR repeats in CEN15. **e**. The numbers of monomer patterns in CEN15 R1L11. **f**. Structure and annotation of CEN1. **g**. The number of monomer patterns in CEN1 R2L6 and R3L6. **h**. Structure and annotation of CEN8. **i**. Dot plots for different HORs in CEN8. D11Z1, D15Z3, D1Z7 and D8Z2 are previously reported HORs. MP is the monomer pattern. # means the number of.

In CEN1, 8, 9, 10, 13 and 19, previously reported HORs were not ranked first but among the top five HiCAT results (Additional file 3: Table S2) due to repeat expansion (Fig. 3 and Additional file 1: Fig. S3). For CEN1, the first HiCAT HOR was R1L2 with two monomers, which was consistent with previously reported dimeric expansions in D1Z7[13] (the HOR name in previous studies was displayed as “D + chromosome number + Z + sequential number”[3, 11]). In the CHM13 genome, a 1.7-Mb inversion in the CEN1 active alpha satellite array[11] split the reported D1Z7 into two HORs, R2L6 and R3L6 (Fig. 3f), with reversed monomer patterns (3-4-7-5-1-2 and 2-1-5-7-4-3, respectively) (Fig. 3g). In R2L6 and R3L6, we also detected expansion of two monomers (3 and 4), and most of them expanded four times (Fig. 3g). The HORs in CEN8 showed location bias. We detected four frequent HORs, namely, R1L15, R2L7, R3L8 and R4L11 (Fig. 3h, i), of which R4L11 was consistent with the reported HOR D8Z2 with 11 monomers[3]. We found that different HORs had different locations in CEN8. R4L11 was mainly distributed in the marginal area, while R2L7 was enriched in the centre. R1L15 and R3L8 were distributed between R4L11 and R2L7.

### Substantial differences between HiCAT and semi-manual HOR annotations in CEN5 and CEN17

Previous studies have reported that CEN1, 5 and 19 contain shared HORs with six monomers (D1Z7, D5Z2 and D19Z3) belonging to supra-chromosomal family 1 (SF1) and are organized as alternating dimers of J1 and J2 monomers[3]. D1Z7 and D19Z3 were detected in CEN1 (R2L6 and R3L6) and CEN19 (R2L6), respectively (Fig. 3f and Additional file 1: Fig. S3a), while D5Z2 was not detected in the top five HiCAT results in CEN5 (Additional file 1: Fig. S4a). The top pattern in CEN5 was R1L12, in which the number of nested units was approximately three times greater than that of canonical units (Fig. 4a). We found that monomer patterns with lengths of 12, 16 and 20 were the three most frequent types of patterns, and the specific pattern was 3-4-6-1-8-5-3-2-(3-4-6-7)×n with n=1, n=2, and n=3, respectively (Fig. 4b, Additional file 1: Fig. S4b). Three patterns were distributed in CEN5 without significant location bias (Fig. 4d).

**Fig. 4.**
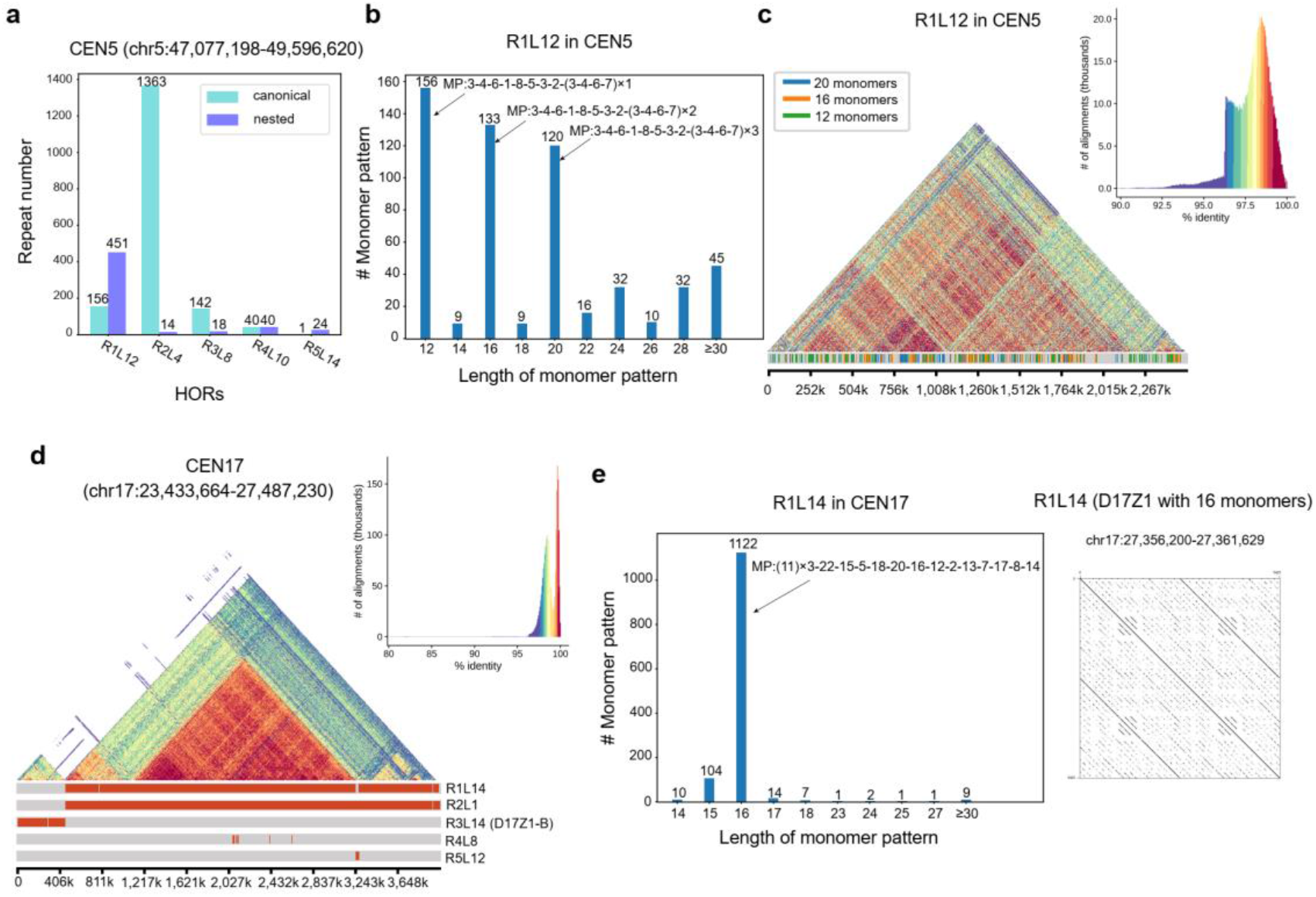
Resolving centromere structure in CHM13 CEN5 and 17. **a**. The HOR repeat number in CEN5. b. The number of monomer patterns in CEN5 R1L12. **c**. Structure and annotation of CEN5 for R1L12 with different monomer pattern lengths. **d**. Structure and annotation of CEN17. **e**. The number of monomer patterns in CEN17 R1L14 and dot plot for R1L14 (D17Z1) with 16 monomers. D17Z1 and D17Z1-B are previously reported HORs in CEN17. MP is the monomer pattern. # means the number of.

Two HORs, D17Z1-B and D17Z1, were reported in CEN17. D17Z1-B with 14 monomers was detected as R3L14 by HiCAT (Fig. 4d), while D17Z1 with 16 monomers was detected as a special case of R1L14 by HiCAT (Fig. 4e). For R1L14, 1,272 HOR units were nested with local TRs, while as few as 10 units were canonical (Additional file 1: Fig. S4c). The monomer pattern was (11)×n-22-15-5-18-20-16-12-2-13-7-17-8-14, and most of the units contained 16 monomers with n=3 (Fig. 4e), consistent with previous reports that D17Z1 belongs to SF3 and experienced triplication of one monomer, e.g., monomer 11 in R1L14 (Additional file 1: Fig. S4d)[17]. Moreover, we also detected other rarer fine structures of R1L14 with different numbers of monomer 11 repeats (Additional file 1: Fig. S4e).

### Comparison with HORmon annotation

We also compared the HORs detected by HiCAT and HORmon[14]. First, we evaluated centromere annotation coverage and continuity in all CHM13 centromeres (Additional file 4: Table S3) and found that the median coverage of both methods was greater than 98% (Additional file 1: Fig. S5a). Moreover, we found that HiCAT significantly outperformed HORmon (*p*-value = 4.6e-7, Wilcoxon rank sum test) in terms of continuity, with fewer annotation breakpoints because the LN-HORs were well captured by HTRM (Fig. 5a and Additional file 1: Fig. S5b).

**Fig. 5.**
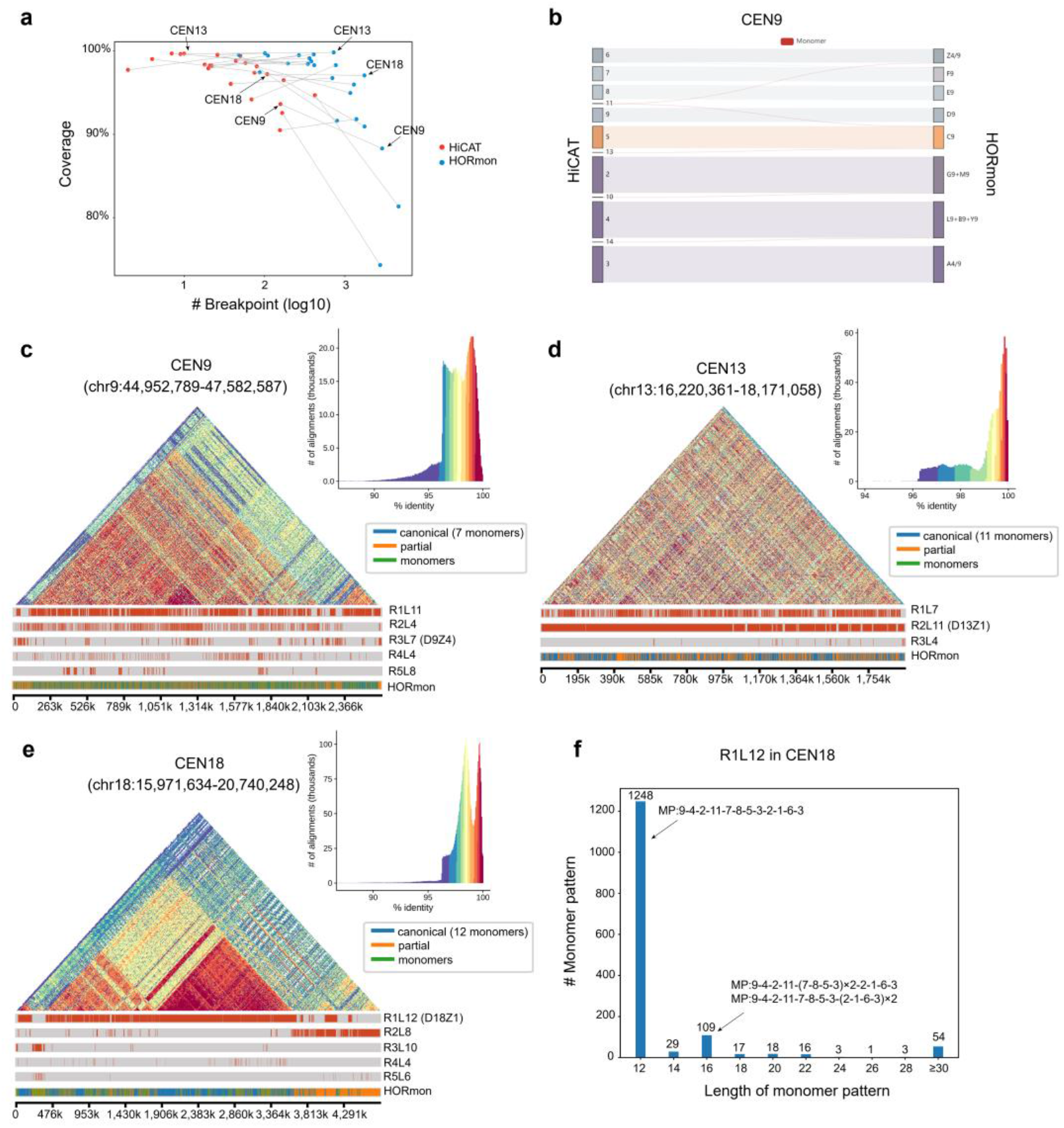
Comparison of HOR annotations between HiCAT and HORmon. **a**. Comparison of the annotation results in terms of coverage and continuity for all CHM13 centromeres. The line between points links the same centromere annotated by HiCAT and HORmon. **b**. Monomer Sankey plot for CEN9 showing the high consistency between the two methods. To display the frequent monomers, we filtered the links with fewer than 10 matches. The complete Sankey plots are shown in Additional file 1: Fig. S5h-j. **c-e**. Structure and annotation of CEN9 (c), CEN13 (d), and CEN18 (e) with two methods. Here, “canonical” represents canonical HORs, “partial” represents partial HORs, and “monomers” represents monomers that do not belong to any HORs. **f**. The number of monomer patterns in CEN18 R1L12. D9Z4, D13Z1 and D18Z1 are previously reported HORs. MP is the monomer pattern. # means the number of.

Next, we further compared HiCAT with HORmon in detail by examining CEN9, 13 and 18, which have extensive LN-HORs. Overall, the monomers inferred by the two methods were largely consistent (Fig. 5b and Additional file 1: Fig. S5c, d). For example, the frequent monomers inferred by HiCAT and HORmon were consistent in CEN9 (Fig. 5b) but different in CEN13 monomer 1 and CEN18 monomers 2 and 3 due to a few-nucleotide difference (Additional file 1: Fig. S5c, d)[14].

For HOR detection, HORmon detected canonical HORs with a monomer pattern as A4/9-(L9+B9+Y9)-C9-D9-E9-Z4/9-(G9+M9) in CEN9[14]. However, monomer F9 with a frequency of 1,193 was annotated as a single monomer in HORmon not belonging to any HORs, reducing the coverage of HOR annotation. In HiCAT, due to the HTRM method, monomer 7 (corresponding to monomer F9 in HORmon) was annotated as a subcomponent of R1L11 with a monomer pattern of (2-3-4-5)×m-9-8-6-(2-3-4-7)×n (Fig. 5c), resulting in an increase in coverage from 88% (in HORmon) to 94% (in HiCAT) (Additional file 1: Fig. S3e). In CEN13 and CEN18, the monomer patterns of HORs were consistent between HORmon and HiCAT; e.g., D13Z1 (HORmon) equalled R2L11 (HiCAT) in CEN13, and D18Z1 (HORmon) equalled R1L12 (HiCAT) in CEN18 (Fig. 5d, e). However, nearly half of the regions were defined as partial HORs or single monomers by HORmon in CEN13 and CEN18 (Additional file 1: Fig. S5e), generating 726 and 1,750 breakpoints, respectively, more than 10 times the number in HiCAT (Additional file 4: Table S3). We reported more fine structures of HORs than HORmon. For example, the canonical monomer pattern R1L12 in CEN18 was 9-4-2-11-7-8-5-3-2-1-6-3, and most of the nested units contained 16 monomers with two expanded parts, 7-8-5-3 or 2-1-6-3 (Fig. 5f, Additional file 1: Fig. S5f). Interestingly, we found that the HOR R2L8 in CEN18 with monomer pattern 9-4-2-11-7-8-5-3 was mainly concentrated on the right end of CEN18, reported as partial HORs in the HORmon annotation (Fig. 5e, Additional file 1: Fig. S5g).

### Annotation of centromere structures in the plant genome

To demonstrate generalization of HiCAT, we applied it to *Arabidopsis thaliana* Col-CEN centromeres assembled by Naish et al.[8]. We first evaluated the accuracy of HOR annotation by comparing our results with the reported representative HOR region of chr2:4,808,994-4,826,785[8]. HiCAT detected this HOR as R18L8 (chr2:4,800,609-4,844,007) with a canonical monomer pattern of 6-4-5-2-6-5-2-4 (Fig. 6a, b, Additional file 1: Fig. S6, Additional file 5: Table S4). Next, we applied HiCAT to all centromeres in the Col-CEN assembly (Additional file 2: Table S1, Additional file 4: Table S4). In contrast to human centromeres, in which most HORs evolved from dimers or pentamers[17], we found one monomer expansion (monomic expansion) in all *Arabidopsis thaliana* centromeres (Fig. 6c, Additional file 1: Fig. S7). For example, in CEN1, the top HOR was R1L2 with canonical pattern 35-4 (Fig. 6c, d), and monomers 35 and 4 experienced a substantial number of expansions (Fig. 6e).

**Fig. 6.**
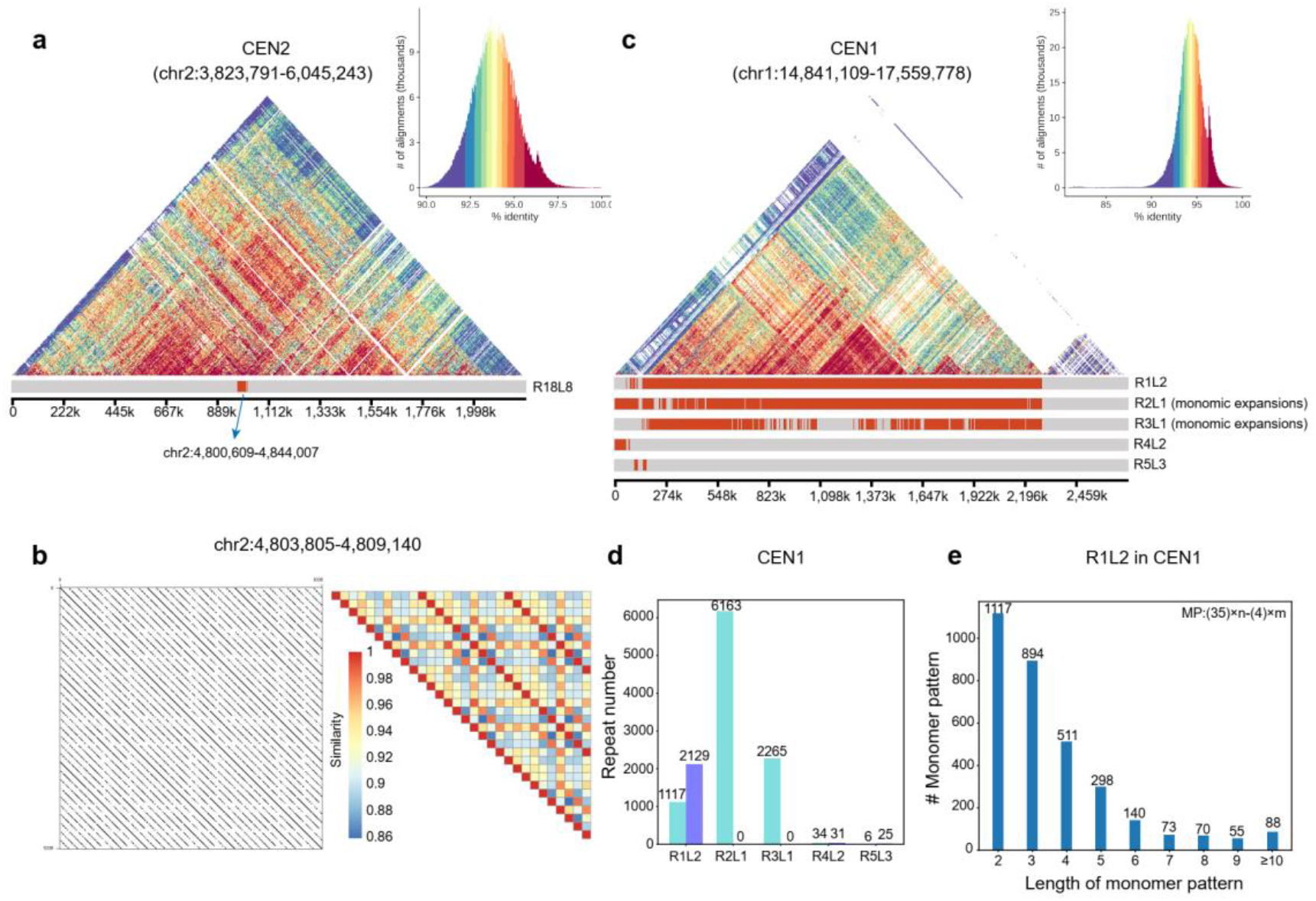
Annotation of centromere structures in *Arabidopsis thaliana* CEN2 and CEN1. **a**. Structure and annotation of CEN2 R18L8. **b**. Dot plot and similarity heatmap for a part of R18L8. The complete dot plot and similarity heatmap are shown in Additional file 1: Fig. S6. **c**. Structure and annotation of CEN1. **d**. The HOR repeat number in CEN1. **e**. The number of monomer patterns in CEN1 R1L2. MP is a monomer pattern. # means the number of.

## Discussion

High-precision and long-read sequencing technologies have revolutionized genome assembly, unlocking complex centromere regions and signalling a new stage in genomics research. The new computing problems introduced by these advances, such as the centromere annotation problem, require novel bioinformatics methods. Here, we propose HiCAT, a generalized computational tool based on the HTRM method and a TR coverage maximization strategy to automatically process centromere annotations. HiCAT is able to correctly annotate HORs and detect fine structures in both human and plant centromeres, especially those with complex LN-HORs. With the emergence of a large number of high-quality genomes, HiCAT will promote the study of pan-species centromere diversity and genetic diseases due to defects in centromeres.

The efficiency of any computational approach is vital for its success. We ran HiCAT on a Linux machine with 28 cores (Intel(R) Xeon(R) Gold 6132 CPU @ 2.60 GHz). In all our tests, the maximum runtime was approximately 2 hours for *Arabidopsis thaliana* CEN5, with a length of 2.8 Mb, and the minimum runtime was only 28 seconds for CHM13 CEN21, with a length of 331 Kb (Additional file 6: Table S5).

As promising as HiCAT is, there are still some technical limitations and future work that we plan to address. The first is the parameter of minimum similarity. Although we applied a TR coverage maximization strategy to guide selection of the similarity threshold, some concerns still should be discussed. If the minimum similarity threshold is set too low, some monomers may be merged, and we may obtain the ancestral state. If the parameter is set too high, the similarity judgement between blocks will be too strict, resulting in too many monomers and leading to failure of HOR detection. In our research, based on a previous study in human centromeres, we set the minimum similarity threshold as 94% since the similarity between HOR units was reported to be in the range of 95-100% in humans[5]. For future newly assembled genomes, this parameter may need to be adjusted to adequately reflect centromere evolution. Although HOR detection from a fully assembled genome gives us comprehensive centromere structures, generating a full genome assembly is still a challenging problem. Annotation of HORs from raw reads is one possible way to obtain and validate centromere structures, and the method named Alpha-CENTAURI has been proposed and applied[18, 19]. We will update HiCAT to accept raw reads as input to extend its application scenarios. Finally, hybrid monomers, which are a concatenate of two or even more monomers, are also important for comprehensively studying centromere architecture and evolution. Hybrid monomers were hypothesized as the “birth” of new frequent monomers and were reported in human CEN5 and CEN8[14]. Currently, HiCAT defines monomers based on only the community detection algorithm, and we will update the monomer inference step to detect hybrid monomers in the future.

## Conclusions

We have presented a generalized computational tool, HiCAT based on the HTRM method and a TR coverage maximization strategy to automatically process centromere annotations. In human and *Arabidopsis thaliana* centromeres, we showed that HiCAT annotation not only were generally consistent with previous inferences but also greatly improved annotation continuity and revealed additional fine structures, demonstrating HiCAT’s performance and general applicability. We believe that with the emergence of a substantial number of high-quality genomes, HiCAT will promote the study of pan-species centromere diversity and genetic diseases due to defects in centromeres.

## Methods

### Datasets in humans and *Arabidopsis thaliana*

We obtained active alpha satellite arrays from the complete sequence of the human CHM13 cell line assembled by the T2T Consortium (version 1.0)[11, 14]. HORmon annotation of CHM13 centromeres was downloaded from https://figshare.com/articles/dataset/HORmon/16755097/2 [14]. We used the Col-CEN assembly of the *Arabidopsis thaliana* genome and obtained the corresponding centromere coordinates from Naish et al.[8]. The centromere regions in both CHM13 and Col-CEN are summarized in Additional file 2: Table S1.

### Generation of the block list and similarity matrix

The first step of HiCAT was to decompose the centromere DNA sequence into the block list based on the input monomer template by StringDecomposer[4] (Fig. 2a). We defined the similarity between blocks *b*_1_ and *b*_2_ as:

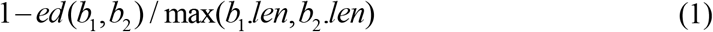

where *ed* is edit distance between *b*_1_ and *b*_2_. *b*.*len* is the block length. We calculated the similarity of each block pair to obtain the similarity matrix. Then, we merged the identical blocks (similarity = 100%) to obtain the merged similarity matrix for improving computing efficiency in the HOR mining step.

### Mining HORs

Based on the merged similarity matrix, we first defined the block graph, whose nodes are blocks and edges are links between any block pairs if their similarity is greater than a given similarity threshold. A series of block graphs were constructed based on the similarity threshold iteratively increasing from the minimum value (by default 94%) to nearly 100% with a specific step (by default 0.5%). Then, we applied the Louvain algorithm[15, 16] to detect communities in each graph and considered each detected community as a monomer. We assigned a unique number to each monomer as its ID. Next, we transformed the block list into monomer sequences based on block communities (Fig. 2b). Since local nested TRs hinder the detection of HORs, we proposed the HTRM method to iteratively detect TRs in monomer sequences. HTRM includes monomer tandem detection, region checking and sequence updating modules. The input of HTRM is a monomer sequence with an upper bound for the length of the TR unit (by default 40 for improving efficiency). We defined a top layer data structure to record non-overlapping TRs with maximum coverage. First, HTRM applied a monomer TR detection module (Additional file 1: Fig. S2a and Additional file 1: Supplementary method) to detect new TRs with a given TR unit length. The initial TR unit length is one. In the second step, we performed region checking (Additional file 1: Fig. S2b) to check for overlap between newly detected TRs (new TRs) and TRs already stored in the top layer (old-TRs). The new TRs and old TRs were modified based on four situations. If there was no overlap between them, the new TRs could be saved in the top layer directly. If partial overlap was detected between old and new TRs, the overlapping new-TRs were removed, and the remaining ones were saved in the top layer. If new TRs covered old TRs, the new TRs replaced old TRs in the top layer. Finally, if new TRs were covered by old TRs, the new TRs were discarded. In the sequence updating module, if the top layer was not updated in the region checking step, the TR unit length for detection was increased by one to redetect TRs. Otherwise, the monomer sequences of the newly saved TR region were compressed. After compression, we redetected the TRs by resetting the TR unit length to one. The details and pseudocode of HTRM are shown in the Additional file 1: Supplementary method. After HTRM, all detected TRs are reported, and their units are normalized; e.g., units of 4-1-2-3, 3-4-1-2 and 2-3-4-1 will be normalized as 1-2-3-4. Then, we merged TRs with the same ordered set of normalized units as a HOR. We calculated the associated HOR coverage of each similarity threshold and chose the threshold with the largest coverage for defining HiCAT HORs. Finally, we ranked HiCAT HORs by HOR score combining the coverage and the degree of local nesting. The HOR score is defined as:

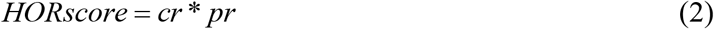

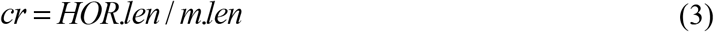

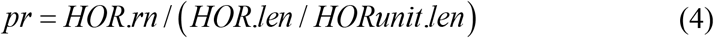

where *cr* is the coverage for the HOR in the input monomer sequence. *pr* represents the degree of local nesting. *HOR*.*len* is the length of the HOR region in the monomer pattern, and *m*.*len* is the length of the monomer sequence. *HOR*.*rn* is the repeat number for the HOR, and *HORunit*.*len* is the length of the HOR unit in the monomer pattern. If the HOR is over-compressed, which means that it contains only a small number of repeats but with high coverage, *HOR*.*rn* will be significantly smaller than *HOR*.*len* / *HORunit*.*len*, and *pr* will balance the coverage and nested degree of the HOR. We named each HOR in each chromosome as “R + (ranking) + L + (*HORunit*.*len*)”. For example, in human CEN11, the first HOR is R1L5.

### Annotation visualization

StainedGlass[20] was used to visualize the TR structures with identity heatmaps, and the window size was set to 2000. We used Gepard[21] to create dot plots. For HiCAT results, within each centromere, we visualized the top five HORs with repeat numbers greater than 10 and reported all detected HORs in the output files.

## Abbreviations

HiCAT: hierarchical centromere annotation tool
CHM: complete hydatidiform mole
T2T: Telomere-to-Telomere
TR: tandem repeat
HOR: higher order repeat
CEN: centromere
HiFi: high-fidelity
SD: StringDecomposer
CE postulate: centromere evolution postulate
LN-HOR: local nested higher order repeat
HTRM: hierarchical tandem repeat mining

## Ethics approval and consent to participate

Not applicable.

## Consent for publication

Not applicable.

## Availability of data and materials

Datasets used for the analyses in this study are summarized in Additional file 3: Table S2. The source code of HiCAT and all annotation results are publicly available at https://github.com/xjtu-omics/HiCAT.

## Authors’ contributions

KY and XY conceived the study. SG, BW and XZ analysed the data. SG and XY developed the program. SG and XY wrote the manuscript. SG completed figures of manuscript. All authors read and approved the final manuscript.

## Acknowledgements

The authors thank Yujing Liu and Peng Jia for their important suggestions and feedback.

