## Additional file 1 for "HiCAT: A tool for automatic annotation of centromere structure"

Supplementary method: Pseudocode for hierarchical tandem repeated mining

Local nested tandem repeats in centromeres hinder the detection of complete continuous high-order repeats. We proposed hierarchical tandem repeated mining (HTRM) method to handle this problem. HTRM recursively detected and compressed local tandem repeats (TRs) in monomer sequence until no tandem repeat is identified.

HTRM takes the input of a monomer sequence (om_seq, original monomer sequence) and an upper bound for the length of repeat unit (UB, by default 40). HTRM defines top_layer to record non-overlap TRs with maximum coverage, all_layer to record TRs in all iterations and tmp_layer to record non-overlap TRs in current iteration that saved into top_layer. In each iteration, HTRM first preforms monomer tandem detection module (Fig. S2a) to detect TRs with a TR unit length (tandem unit length, tul, starts with one). In monomer tandem repeat detector, we first defined candidate pattern (cp) to record index in monomer sequence with same monomer and then using the distance database (d_db) to record two same monomers that their distance equals to tul. In d_db, we recorded the current and previous monomer positions (current start and preivious start, cs and ps). Finally, we sorted the d_db based on the ps and detecting continuous region in sort_d_db as TRs.

After monomer tandem detection module, if there is no TR detected, tul plus one and re-detected. If tul exceeds the minimum value between the length of current monomer sequence and UB, then stop iteration and return TRs in all_layer. If TRs are detected, HTRM preforms region checking module (Fig. S2b) to check the overlap between new TRs and TRs in top_layer. Since the monomer sequence in each iteration will be compressed, HTRM defines cm_seq_index (current monomer sequence index) to map the index in current monomer sequence (cm_seq) to the index in input original monomer sequence (om_seq) and then checking the overlap using the index in original monomer sequence (os and oe, start and end in original monomer sequence). There are four situations. The first one is that new TR not overlap with all TRs in top_layer. HTRM add new TR to top_layer, tmp_layer and all_layer. The second one is that new TR partially overlaps with some TRs in top_layer. The new TR needs to be modified the start and end (mos and moe) to not overlap with old TRs in top_layer then adding to top_layer, tmp_layer and all_layer. The third one is new TR covered some TRs in top_layer. Then, these TRs in top_layer needs to remove and added new TR to top_layer, tmp_layer and all_layer. Finally, new TR can be a part of some TRs in top_layer. In this situation, this new TR needs to be abandoned. If there are no item in tmp_layer, tul needs to plus one and re-detected TRs. After the region checking module, HTRM preforms sequence update module to compress current monomer sequence and cm_seq_index based on tmp_layer and tul will be set to one for next iteration.

The pseudo code of HTRM is as follows. And the source code is in <https://github.com/xjtu-omics/HiCAT>

**Algorithm 1: Hierarchical tandem repeated mining**

**HTRM**(om_seq,UB):

nm_seq = om_seq

nm_seq_index mapping monomer index to om_seq

top_layer record non-overlap tandem repeats with maximum coverage

all_layer record tandem repeats in all iterations

tul = 1

**while** tul < min(UB,nm_seq.length):

cm_seq = nm_seq

cm_seq_index = nm_seq_index

// Monomer tandem detection module

tr_list = MTRD(cm_seq,tul)

// Region checking module

tmp_layer record non-overlap tandem repeats in current iteration that saved into top_layer.

**foreach** r **in** tr_list:

cs = r.start

ce = r.end

os = cm_seq_index[cs]

oe = cm_seq_index[ce]

**if** top_layer is empty:

// info records repeat number and each unit

tri = (cs, ce os, oe, info) top_layer.add(tri)

tmp_layer.add(tri)

all_layer.add(tri)

**continue**

**foreach** tri **in** top_layer:

ts = tri.os

te = tri.oe

**if** os <= te && ts < os && oe > te:

region overlaps with tri to left

**if** os < ts && oe >= ts && oe < te:

region overlaps with tri to right

Modify (cs,ce,os,oe) to (mcs,mce,mos,moe) make sure non-overlap

**foreach** tri **in** top_layer:

ts = tri.os

te = tri.oe

**if** mos <= ts && te <= moe:

region covers tri

remove tri from top_layer

**if** moe < ts || te < mos:

continue

**if** os >= ts && oe <= te:

move to next r

tri = (mcs,mce,mos,moe,info)

top_layer.add(tri)

tmp_layer.add(tri)

all_layer.add(tri)

**if** tmp_layer is empty:

tul += 1

**continue**

// Sequence update module

**foreach** tri **in** tmp_layer:

compress cm_seq and cm_seq_index

nm_seq = compressed cm_seq

nm_seq_index = compressed cm_seq_index

tul = 1

**return** all_layer

**Algorithm 2: Monomer tandem repeat detector**

**TRD**(cm_seq,tul):

cp record index in monomer sequence with same monomer

**foreach** i **in** cm_seq.length:

cp[cm_seq[i]].add(i)

d_db record same monomer that distance equals to tul

**foreach** key **in** cp:

pdb = cp[key]

**foreach** j **in** pdb.length:

cs = pdb[j]

**if** j == 0:

**continue**

cdb = pdb[:j]

**foreach** k **in** reversed(cdb):

ps = k

d = cs – ps

**if** d == tul:

d_db.add([ps,cs])

**if** d > tul:

**break**

sort d_db based on ps obtain sort_d_db

init_tr_list record each continuous region in sort_d_db

**foreach** r **in** tr_list:

**if** int(r.length)/tul) > 1:

r = r[:int(r.length)/tul) * tul]

tr_list.add(r)

**return** tr_list

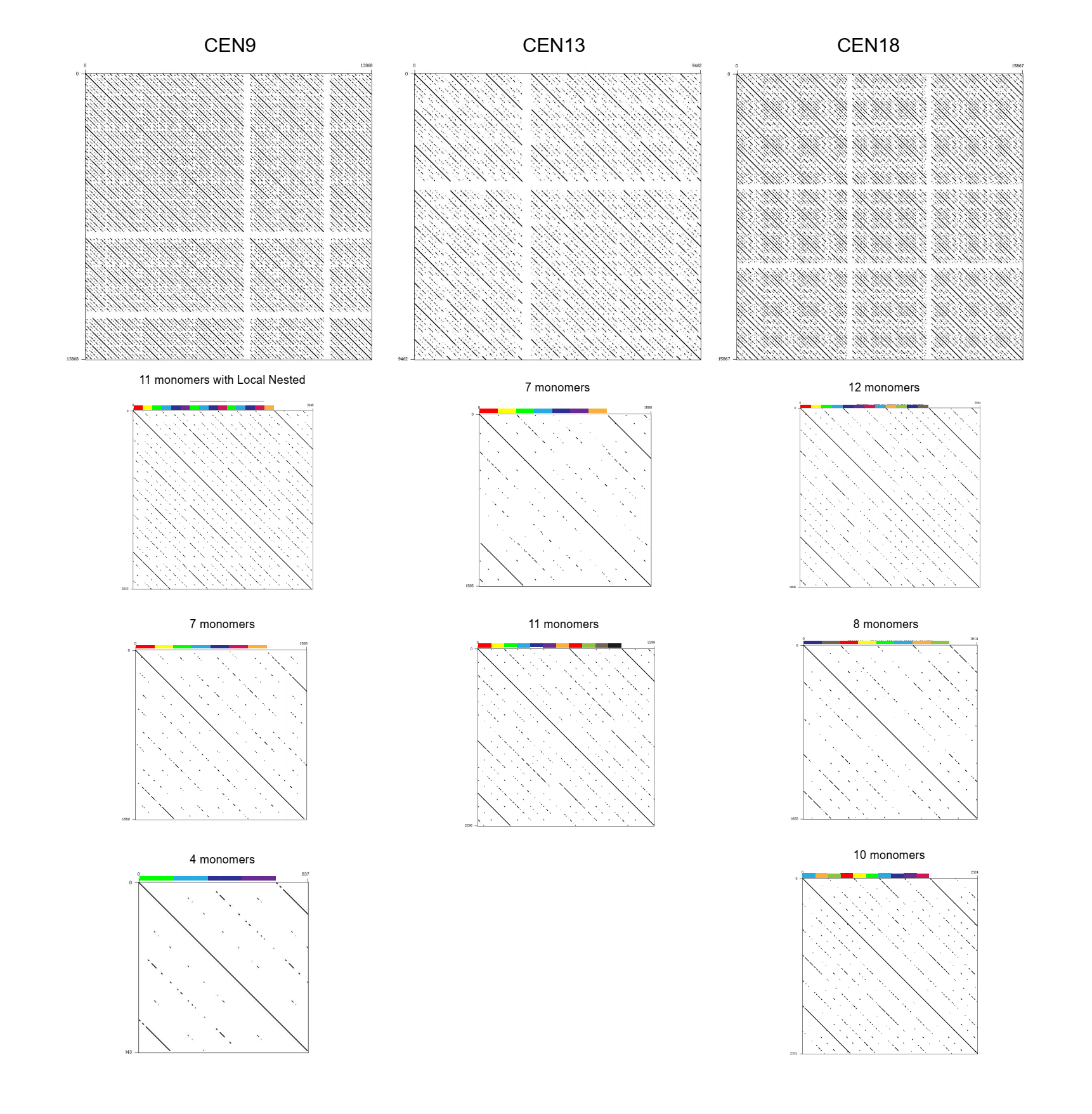

**Supplementary figure S1| Multiple length of HOR units with shared monomers in human CHM13 CEN9, 13 and 18**. Different color rectangles represent different monomers.

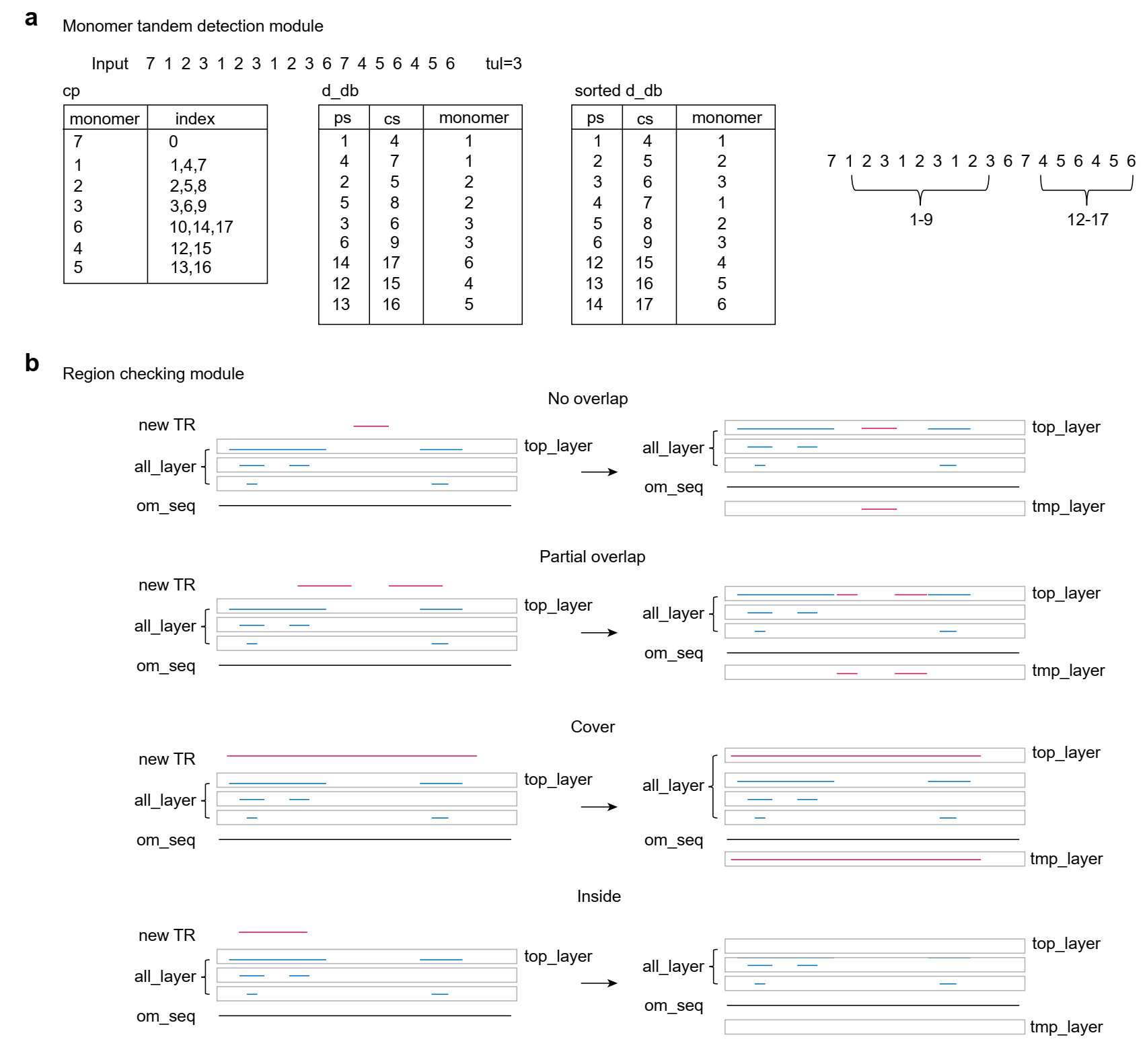

**Supplementary figure S2| Two modules in hierarchical tandem repeated mining method. a.** Monomer tandem detection module. The input are monomer sequence and a fixed unit length (tandem unit length, tul). cp record index in monomer sequence with same monomer. d_db record same monomer that distance equals to tul. cs is current start and ps is previous start. For example, for cs = 4 and ps = 1, ps – cs = 3 equals to tul = 3. sort d_db is sorted d_db based on ps. **b.** Region checking module contains four situations. TR is a tandem repeat. HTRM defines top_layer to record non-overlap tandem repeats with maximum coverage, all_layer to record tandem repeats in all iterations and tmp_layer to record non-overlap tandem repeats in current iteration that saved into top_layer.

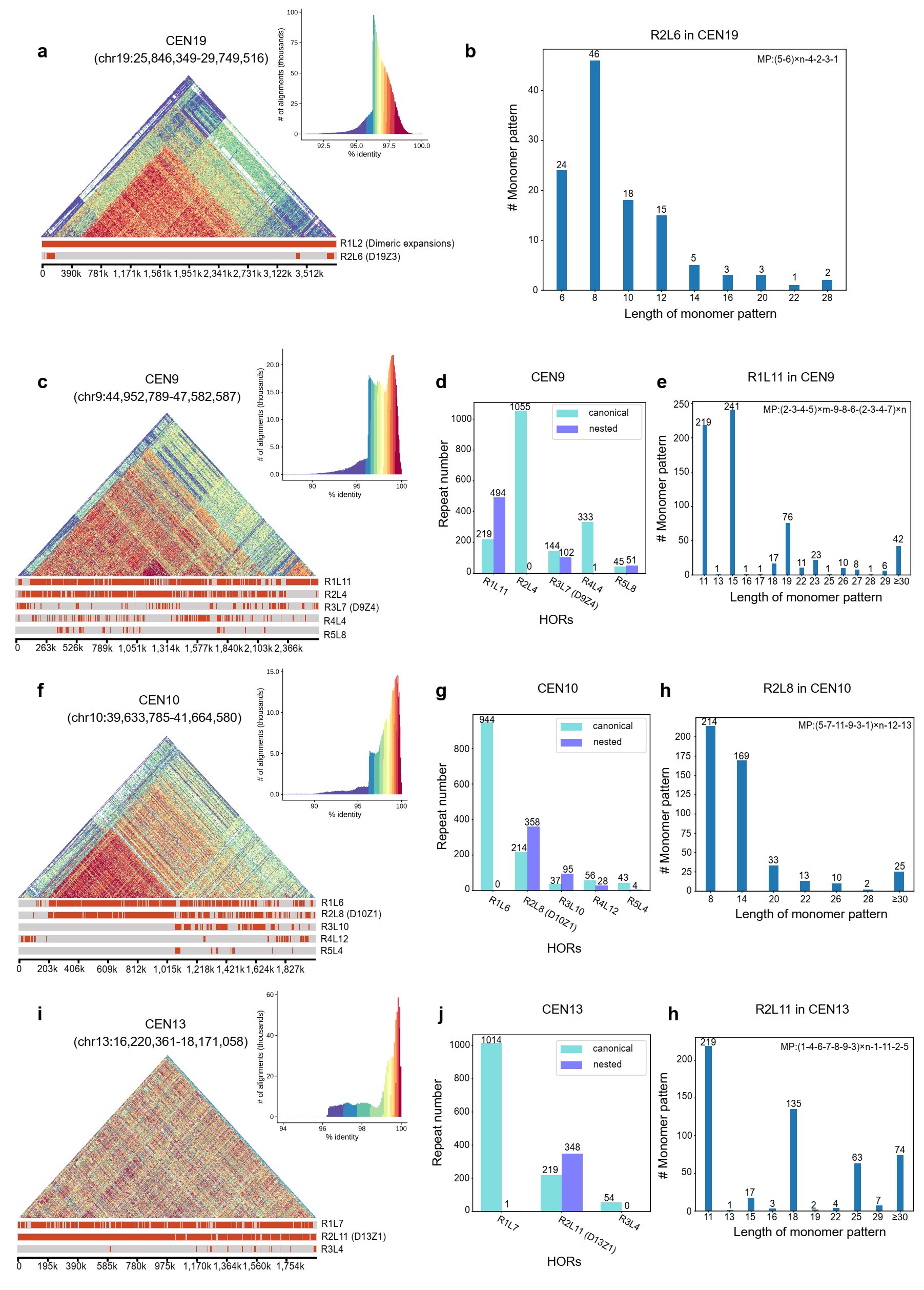

**Supplementary figure S3| The annotation in human CHM13 CEN19, 9, 10 and 13. a.** Structure and annotation of CEN19. **b.** The number of monomer pattern in CEN19 R2L6. **c.** Structure and annotation of CEN9. **d.** The HORs repeat number in CEN9. **e.** The number of monomer pattern in CEN9 R1L11. **f.** Structure and annotation of CEN10. **g.** The HORs repeat number in CEN10. **h.** The number of monomer pattern in CEN10 R2L8. **i.** Structure and annotation of CEN13. **j.** The HORs repeat number in CEN13. **h.** The number of monomer pattern in CEN13 R2L11. D19Z3, D9Z4, D10Z1 and D13Z1 were previous reported HORs. MP is monomer pattern. # means the number of.

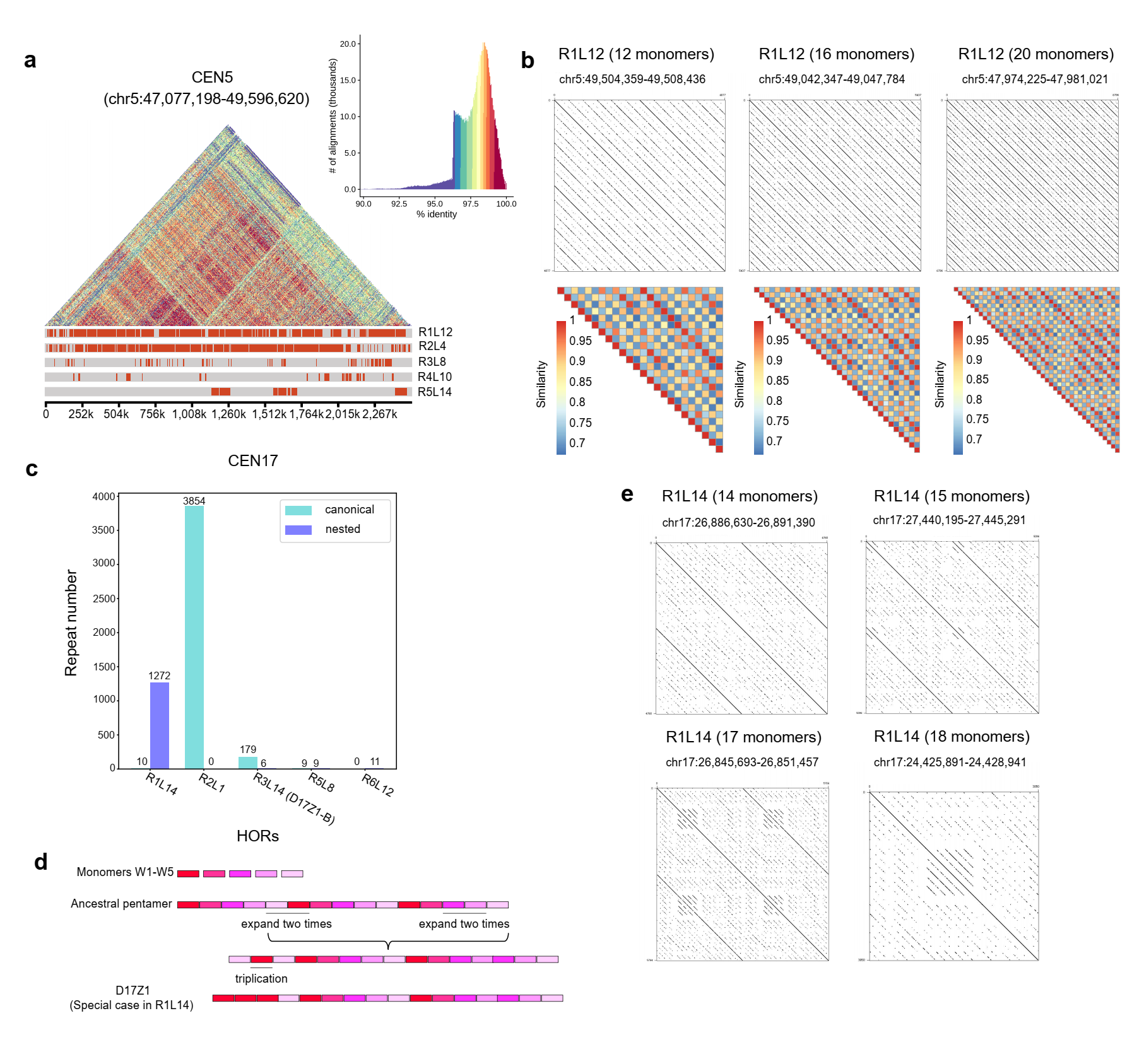

**Supplementary figure S4| The** **annotation in human CHM13 CEN5 and 17. a.** Structure and annotation of CEN15. **b.** Dot plots and similarity heatmaps for R1L12 with different monomer length. **c.** The HORs repeat number in CEN17. **d.** Evolution history of D17Z1 (R1L14 with 16 monomers). **e.** Dot plots for R1L14 with 14, 15, 17 and 18 monomers. D17Z1-B was previous reported HOR in CEN17. MP is monomer pattern. # means the number of.

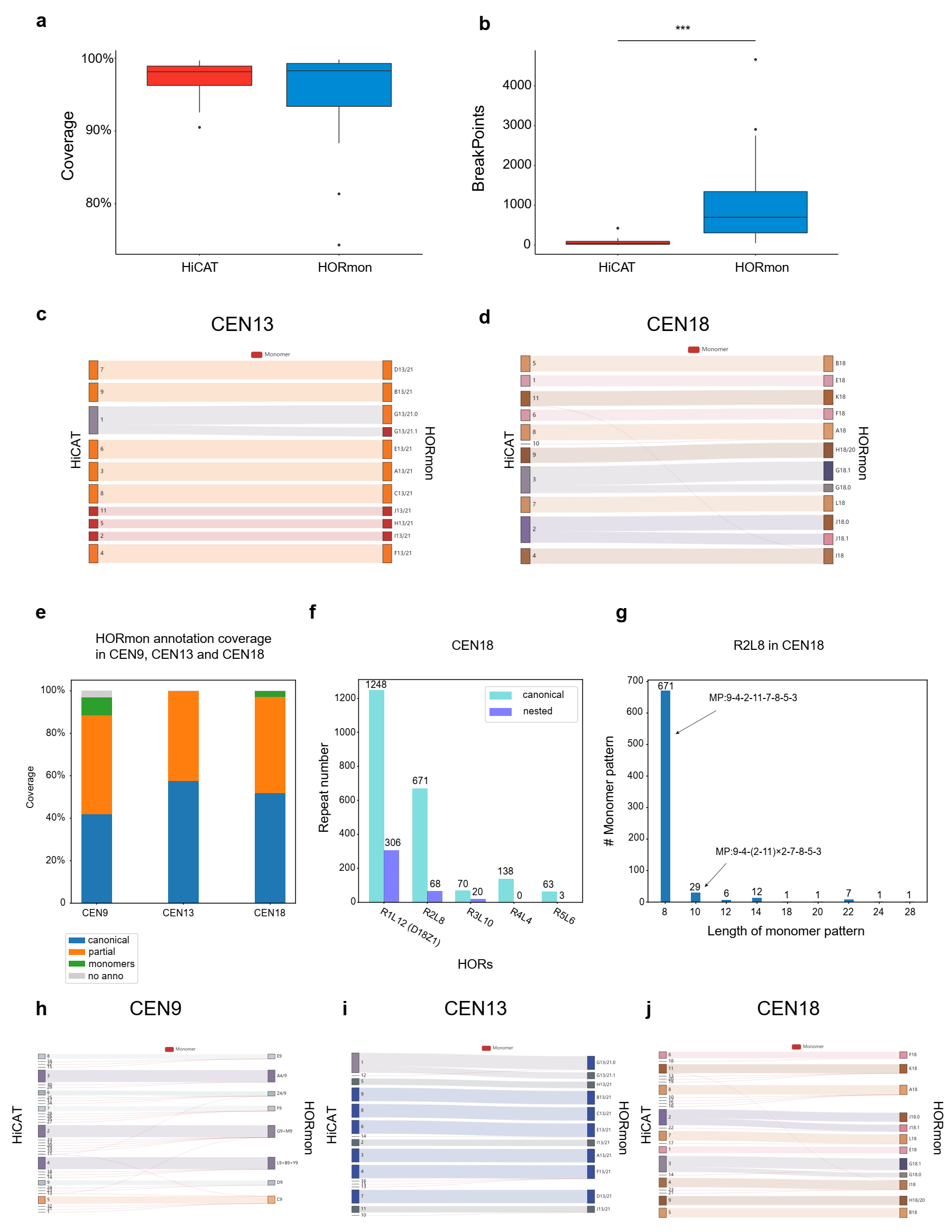

**Supplementary figure S5| Compared with HORmon in human CHM13. a.** Compared with HORmon in annotation coverage. **b.** Compared with HORmon in annotation continuity. *p*-value = 4.6e-7. *** represents *p*-value < 0.001, Wilcoxon rank sum test. **c.** Monomers sankey plot for CEN13. **d.** Monomers sankey plot for CEN18. To display the frequent monomers, we filtered the links with match number less than 10. **e.** HORmon annotation type coverage in CEN9, 13 and 18. “canonical” is represented canonical HORs. “partial” is represented partial HORs and “monomers” is represented monomers not belong to any HORs. **f.** The HOR repeat number in CEN18. **g.** The number of monomer pattern in CEN18 R2L8. **h-j.** Unfiltered monomers sankey plot for CEN9 (h), CEN13 (i) and CEN18 (j). MP is monomer pattern. # means the number of.

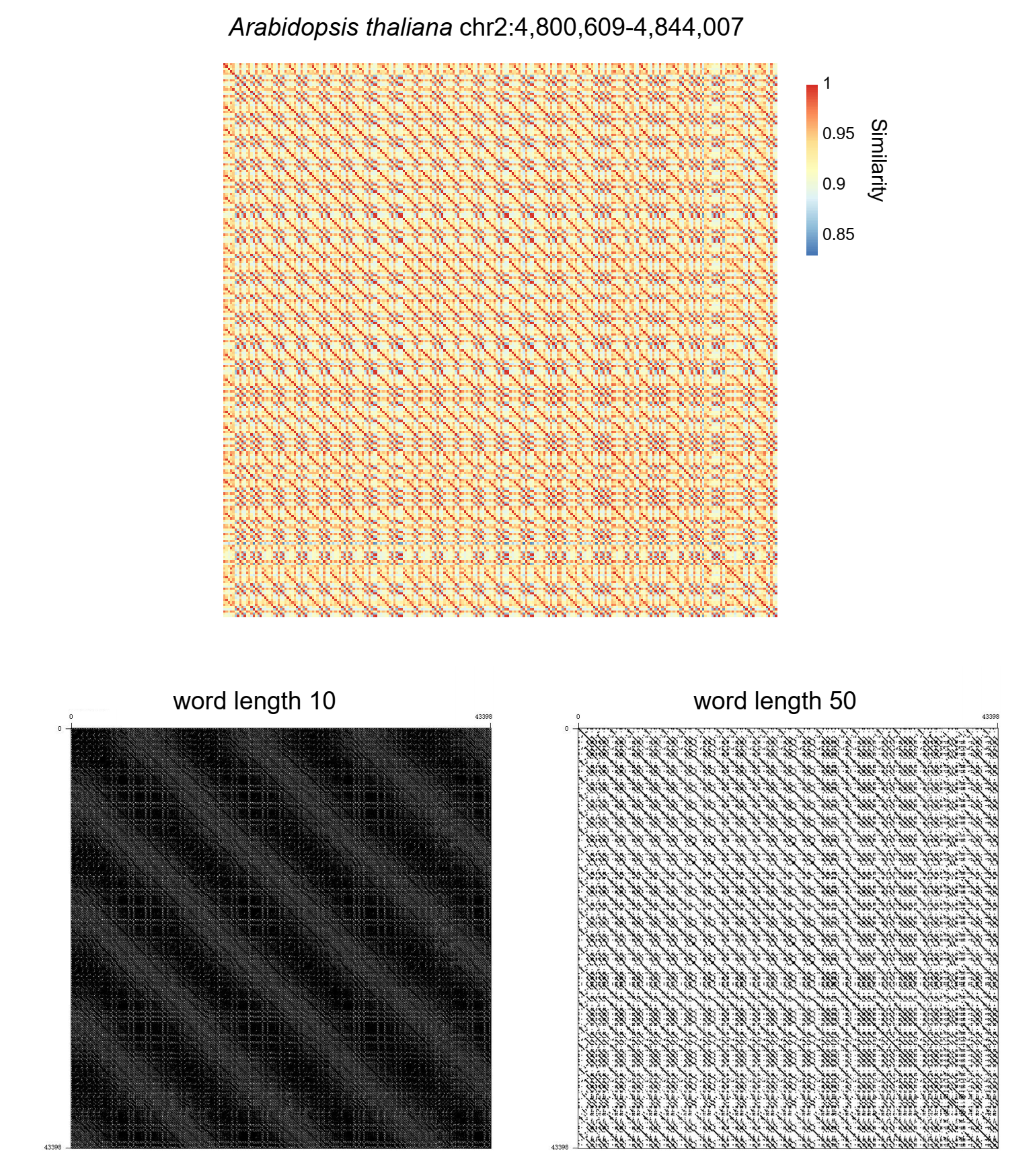

**Supplementary figure S6| Dot plots and similarity heatmap for R18L8 in *Arabidopsis thaliana* CEN2.**

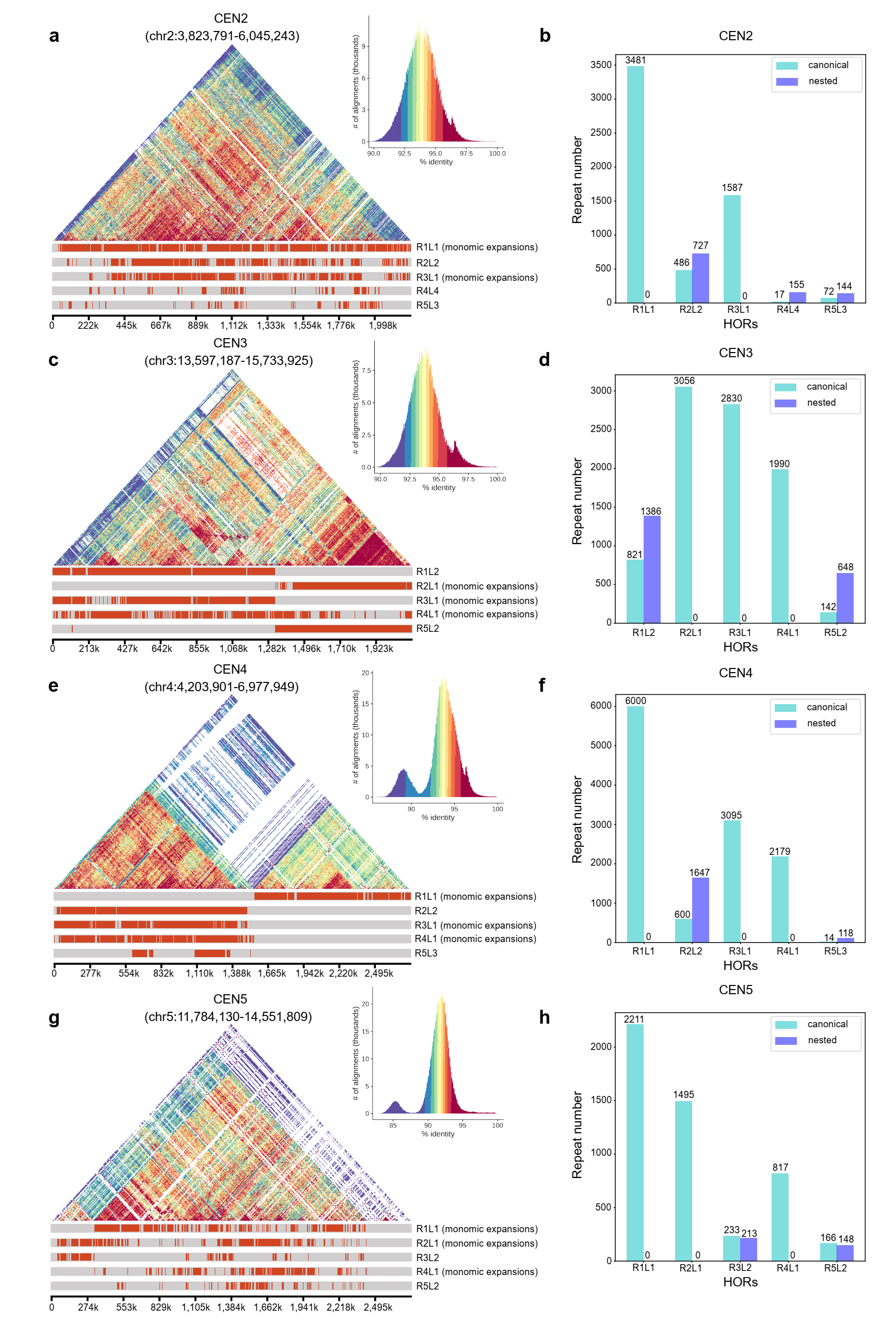

**Supplementary figure S7| Annotation centromere structures in *Arabidopsis thaliana* CEN2, CEN3, CEN4 and CEN5. a.** Structure and annotation of CEN2. **b.** The HOR repeat number in CEN2. **c.** Structure and annotation of CEN3. **d.** The HOR repeat number in CEN3. **e.** Structure and annotation of CEN4. **f.** The HOR repeat number in CEN4. **g.** Structure and annotation of CEN5. **h.** The HOR repeat number in CEN5.
